# Exon-Resolved Dissection of Shared, Unique, and Antagonistic Functions of ZFAS1 Isoforms

**DOI:** 10.64898/2026.05.22.727302

**Authors:** Sébastien Soubeyrand, Paulina Lau, Ruth McPherson

**Author notes:** Corresponding author: Sébastien Soubeyrand (SS).

## Abstract

The long noncoding RNA *ZFAS1* plays a role in cell proliferation and has been linked to cancer development and prognosis. However, the *ZFAS1* locus is predicted to encode over 40 *ZFAS1* splice variants, which remain largely uncharacterized. To shed light on the role of *ZFAS1* in hepatocyte models, we examined the transcriptome-wide effects of separately targeting three exons present in representative *ZFAS1* variants with gapmer antisense oligonucleotides. Evidence of cross-exon compensatory regulation was obtained by qRT-PCR. Although targeting resulted in a subset of concerted transcript perturbations-indicating specificity and shared functionalities-the overall effects were predominantly non-redundant. Overrepresentation analyses revealed that proximal exon and distal exon targeting affected cell cycle and transcription regulation, respectively. Strikingly, interrogation of the entire transcriptome with gene set enrichment analysis identified a shared subset of pathways related to cell-cycle control and translation, which were affected antagonistically in an exon-dependent manner. Whereas targeting the proximal exon was predicted to broadly compromise cell-cycle and translational functions, targeting distal exons produced contrasting effects on these processes. Together, these findings demonstrate that the arrangement of *ZFAS1* exons can markedly modulate its function.

## Introduction

Long non-coding RNAs are a class of RNAs defined by their lack of protein coding potential and length (> 200 nucleotides), exhibiting considerable functional diversity (1,2). Relative to mRNAs, lncRNAs are less conserved, tend to be expressed at lower levels, and undergo increased splicing variability resulting in greater transcript diversity (1,3,4). The lncRNA *ZFAS1* gene encodes a group of transcripts that have been linked to cancer, inflammation, calcium regulation, cell proliferation, and ribosomal biogenesis (5–9). This multiplicity of roles may reflect cell-type properties or variant-specific roles. Indeed, current *ZFAS1* models at Ensembl (Release 15) predict up to 47 distinct transcript variants (**Fig. 1**). Five splice variants similar to *ENST000000371743.8* (*ENST1743*) have been identified in a breast cancer cell line. Little is known about the alternate 3’ exons (e.g., *ENST000000836971.1*) or the alternate *ENST000000618800.1* (*ENST8800*) variant, although we recently demonstrated the presence in primary hepatocytes of a low-abundance transcript spanning *ENST1743* exon 2 *to ENST800* exon 2, similar to *ENST00000652916.1* (10).

**Figure 1.**
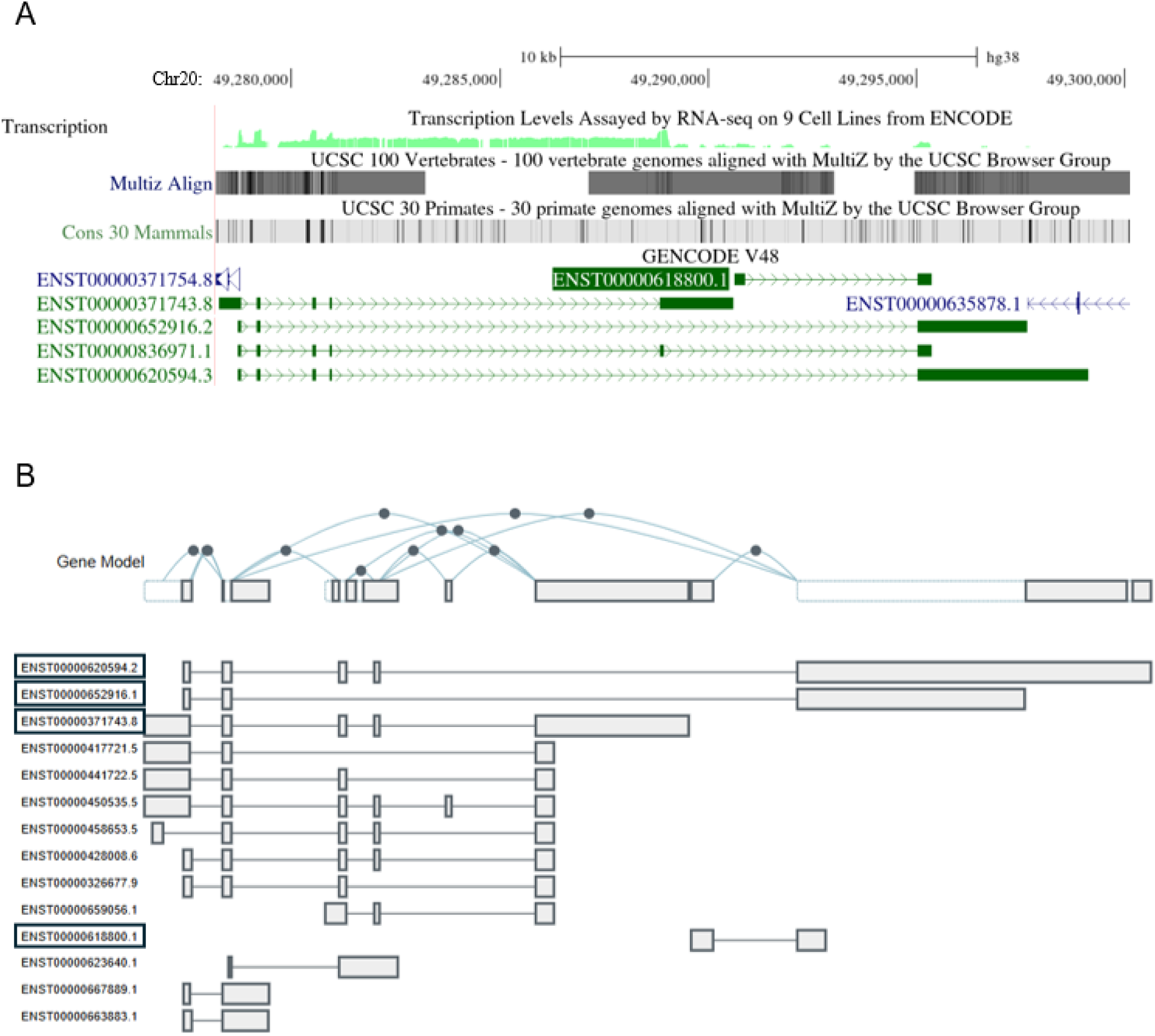
Representative ZFAS1 variants according to GENCODE and GTEx consortia. A, UCSC browser snapshot (https://genome.ucsc.edu/index.html) showing transcription levels in 9 cell lines (top), conservation in primates and vertebrates (middle), and five representative ZFAS1 variants (green). B, Exon arrangement of ZFAS1, as modeled by GTEx (https://www.gtexportal.org/home/gene/ZFAS1/exonExpressionTab). Transcripts with boxed IDs are also displayed in the UCSC browser.

*ZFAS1* expression correlates with a poor cancer prognosis and has been proposed as a blood-derived biomarker for hepatocellular carcinomas (5,8). To better understand the role of *ZFAS1* in liver pathophysiology, we recently characterized *ZFAS1* in various hepatocyte models, wherein we reported an association between *ZFAS1* and the unfolded protein response (UPR) (10). Although the suppression of all *ZFAS1* variants led to modestly reduced cell viability, splice variants harbouring proximal exons (i.e. relative to the promoter) were more responsive to UPR than distal and less abundant forms, alluding to functional diversity. However, *ZFAS1* suppression had limited impact on the UPR response, as assessed by the abundance of pivotal UPR mediators. Here, to clarify the role of *ZFAS1* variants, we have examined the transcriptomic consequences of targeting three representative *ZFAS1* exons in the widely used hepatoblastoma cell line (HepG2 cells). Our findings point to both shared and unique, occasionally antagonistic, contributions of *ZFAS1* variants, underscoring the functional complexity of the *ZFAS1* locus.

## Materials and Methods

### Antisense oligonucleotide transfection

HepG2 cells (a total of 4 distinct passages, 13 to 22) maintained in Low Glucose DMEM (Gibco) were transfected for 48 h with a total of 7 ASOs: 3 pairs of antisense oligonucleotides (ASOs), each pair targeting one of three ZFAS1 exons, or a control ASO (see **Supplementary Sequence Information**). Cells were seeded at 100,000 cells in 500 µL of media per well in 24-well plates and were transfected immediately with 30 pmol of gapmer ASO and 1 µL of RNAiMAX (Invitrogen), prepared in 100 µL of Opti-MEM (20 min incubation).

### RNA purification, cDNA synthesis, and quantitative reverse transcription PCR (qRT-PCR)

Cells were lysed 48 hours after transfection with 200 µL of TriPure reagent (Roche), and the lysate was column-purified (Direct-Zol RNA MiniPrep kit; Zymo Research). The *ZFAS1* qPCR primer pairs had comparable amplification efficiencies (>95%) as determined by serial dilutions of HepG2 cDNA. For a typical reaction *ENST1743* exon 3, *ENST8800* exon 1, and *ENST8800* exon 2 yielded Cp values of ∼22, 31, and 33, respectively. *PPIA*, which we have used as a robust reference transcript over the years, was used for normalization using the ΔΔCt method; according to our array data (GSE308839), *PPIA* abundance was unaffected (p> 0.25) by ASO treatments (–1.1 (ASO3+4), 1.0 (ASO5+6), and 1.1 (ASO6+7) –fold change vs control ASO). Oligonucleotide sequences are detailed in **Supplementary Sequence Information**. RT-PCR was performed using 1 µL of cDNA in 25 µL reactions for 40 cycles with Terra PCR Direct Polymerase (Takara Bio).

### Transcription array analysis

RNA from the 4 sets of replicates analyzed by qRT-PCR was sent to the Centre for Applied Genomics (The Hospital for Sick Children, Toronto, Ontario, Canada) for processing. Biotin-labeled cRNA was hybridized to human Clariom S arrays (Affymetrix). The Clariom S assay was selected because it focuses on well-annotated genes, thus allowing for robust transcriptome ontology analyses. Thus, it excludes by design most lncRNAs, including *ZFAS1*, that remain insufficiently uncharacterized. Results were processed with the Expression Console and analyzed with the Transcriptome Analysis Console (Affymetrix). The data are publicly available at the Gene Expression Omnibus (GSE308839).

### Enrichment analysis

For overrepresentation analyses, gene identifiers present on the Clariom S were used as the reference list (corresponding to 20,435 transcripts with NCBI Entrez IDs). The most significantly impacted (p < 0.05, >|1.5-fold| change) transcript IDs were mapped to Reactome terms using WebGestalt (https://www.webgestalt.org/) (11). Replication was performed using ShinyGO and g:Profiler (12,13). For the g:Profiler only the Entrez gene identifiers with annotations were considered for statistical purposes; the default g:SCS algorithm was used to calculate significance. For Gene Set Enrichment Analysis (GSEA) with WebGestalt, the entire list of Entrez Gene identifiers was used, along with the corresponding linear fold changes. Unless otherwise mentioned, only Reactome and Gene Ontology hits achieving or exceeding FDR significance (<0.05) were retained. REVIGO (http://revigo.irb.hr/) was performed using the default settings (14). Venn diagrams were generated with E Venn (https://www.bic.ac.cn/test/venn/<u>#/</u>) (15).

## Results

### Local effects of *ZFAS1* suppression

*ZFAS1* exons were selected to cover representative *ZFAS1* variants, including *ENST1743* exon 3, *ENST8800* exon 1, and *ENST8800* exon 2 (the latter being shared with additional *ZFAS1* variants sharing exons with *ENST1743*) (**Fig. 2A**). HepG2 cells were treated with three pairs of gapmers, each pair targeting a different exon (or a control gapmer), followed by exon quantification by qRT-PCR. A 48-h treatment regimen was chosen to capture proximal *ZFAS1* contributions and minimize longer-term impacts, which we reasoned may represent more generic adaptive responses to stress. Targeting *ENST1743* exon 3 decreased its abundance by 75%, and was associated with a significant (∼2-fold) upregulation of *ENST8800* exon 2 (**Fig. 2B**). Interestingly, targeting *ENST8800* exon 1 also increased *ENST8800* exon 2 abundance by approximately 16-fold. By contrast, targeting *ENST8800* exon 2 did not alter expression other exons. To complement this exon-centric quantification, we also quantified three inter-exon signals corresponding to transcripts we previously identified, revealing a consistent pattern (**Fig. 2C**) (10).Notably, possibly owing to the very low basal expression of *ENST8800* (see Discussion), targeting its exons did not measurably affect transcript abundance (**Fig. 2D, E**). Nevertheless, inter-exon effects and substantial *ENST1743* suppression supported the potential efficacy of these interventions, warranting further investigation.

**Figure 2.**
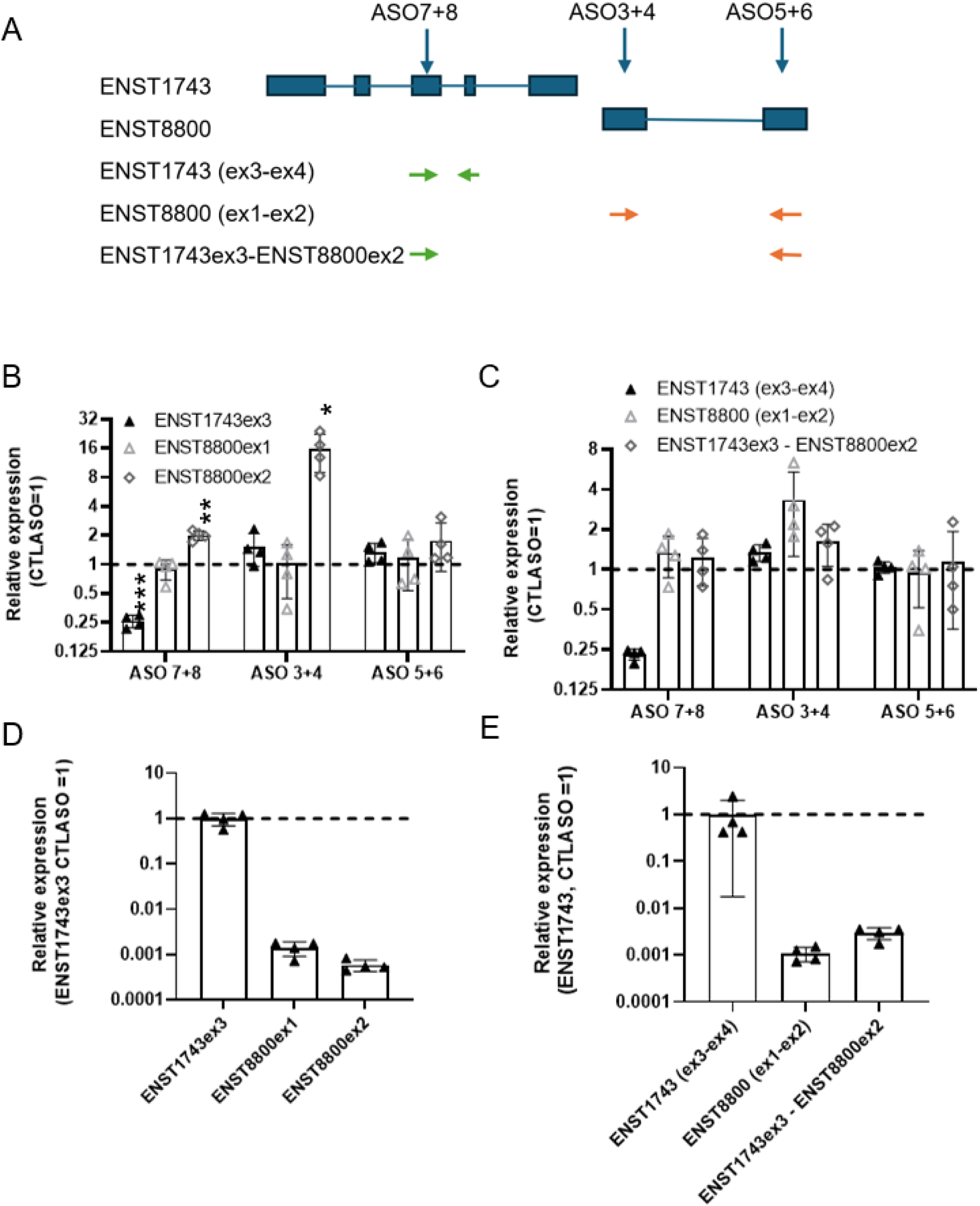
Exon– and transcript-level effects of ZFAS1 suppression. A, schematic of the ASO pairs to suppress ZFAS1. Arrows indicate qPCR primer pairs used to measure inter-exon signals. The *ENST1743* and *ENST8800* correspond to Ensembl transcripts *ENST00000371743.8* and *ENST00000618800.1*, respectively (**Fig. 1)**. B, C, impact on exon-level and trans-exon abundance, respectively. Suppression efficacy measured by qPCR (relative to PPIA, ΔΔCt method) and expressed relative to the CTLASO value. D, E, Intensity of the exon (D) and inter-exon (E) qPCR signals from the CTLASO values (relative to the *ENST1743* exon 3 value)

### Transcriptome-wide effects of *ZFAS1* suppression

The *ZFAS1*-targeted HepG2 cell transcriptomes were then profiled using Clariom S transcription arrays. The Clariom S focuses on well-annotated genes, making it well suited for transcriptome profiling analyses. Targeting *ZFAS1* exons resulted in the altered expression (p < 0.05, >|1.5-fold|) of 1015, 1350, and 351 transcripts for ASO7+8 (*ENST1743* exon 3), ASO3+4 (*ENST8800* exon 1), and ASO5+6 (*ENST8800* exon 2), respectively. A relatively small overlap in affected (p < 0.05, >|1.5-fold| change) transcripts was observed across the suppressions, although the shared groups were enriched (**Fig. 3A).** Importantly, these shared terms were overwhelmingly directionally concordant (**Supplementary Table S1**). For instance, 219/222 and 14/14 hits were directionally concordant for ASO3+4/ASO7+8, and 3-way intercepts, respectively. By comparison, less overlap was observed between the *ENST8800* exons than between *ENST8800* exons and *ENST1743* exon 3, suggesting that hybrid forms may predominate. Nevertheless, most of the hits (87%) were not shared, indicating different (qualitative or quantitative) contributions and/or off-targeting.

**Figure 3.**
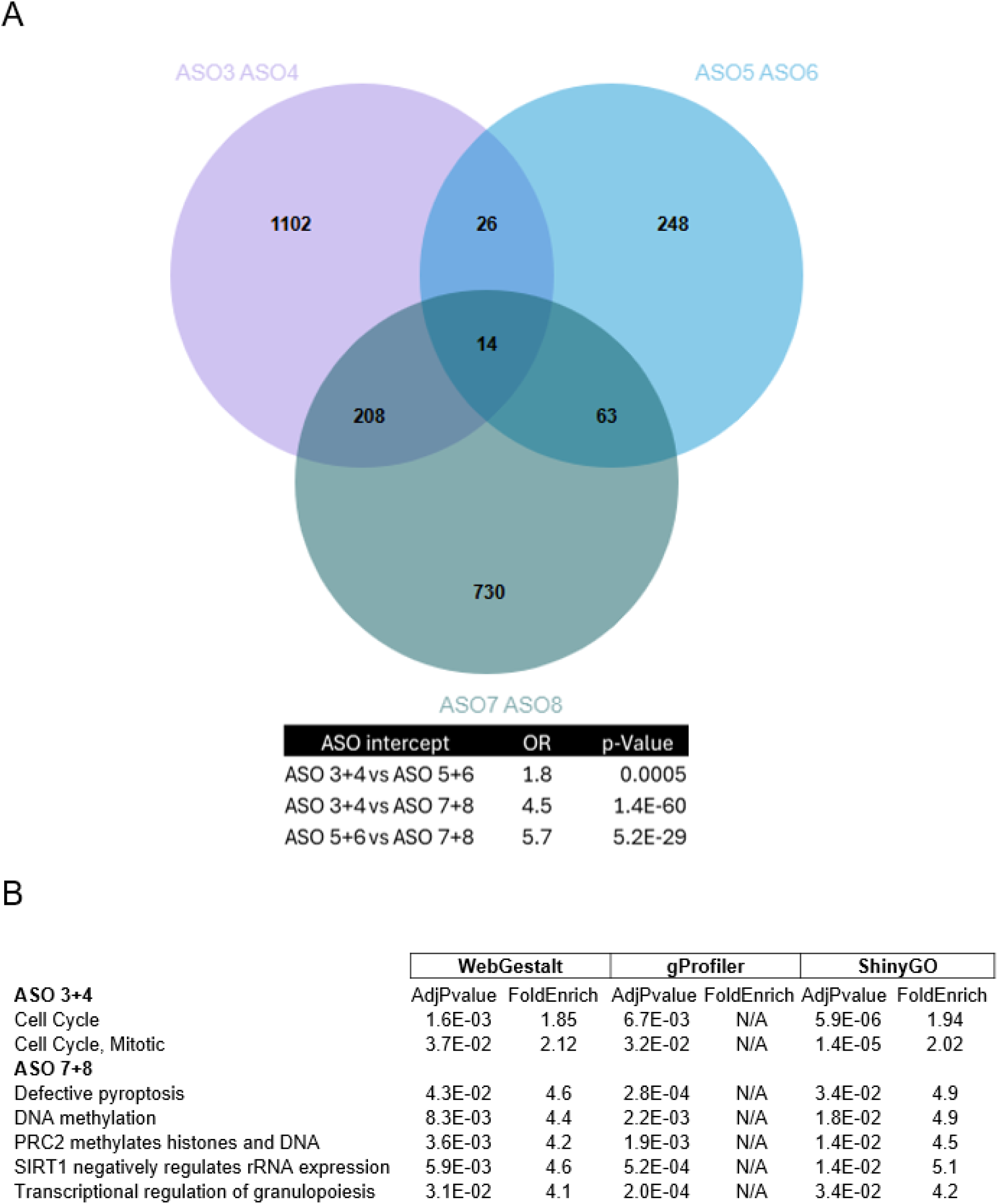
Comparison of the transcriptional profiles of *ZFAS1*-suppressed cells. A, Venn diagram showing the overlap between the perturbed (>|1.5|-fold, p< 0.05) transcripts by the *ZFAS1* ASOs. The table displays the enrichment of the intercept (Fischer’s test; odds ratio (OR), and p-value). B, terms shared by all 3 ORA pipelines. See also **Supplementary Table S2** for a complete list of terms. AdjPvalue: P-value corrected for FDR. FoldEnrich: fold-enrichment in the category. g:Profiler does not provide fold-enrichment information (N/A).

### Overrepresentation analysis of *ZFAS1*-suppressed cells

Next, overrepresentation analysis (ORA) using the WebGestalt pipeline was performed to examine the consequences of individual exon targeting on cell function. The Reactome pathways dataset was selected because of its well-curated and comprehensive content (16). Incorporating the totality of affected (p < 0.05, >|1.5-fold| change) transcripts for each treatment identified a wide range of FDR-significant-enriched Reactome terms for ASO3+4 (71 terms) and ASO7+8 (62 terms) treatments (**Supplementary Table S2**). Interestingly, only one category (“Chromosome maintenance”) was shared by both treatments. Whereas repeating the analysis on the shared populations of significantly affected transcripts was unable to identify FDR-significant pathways, leveraging non-overlapping transcripts to delineate unique contributions identified 79 (ASO3+4) and 51 (ASO7+8) FDR-significant terms which extensively intersected with the corresponding global analyses (**Supplementary Table S2**). Together these results suggest that targeting these three exons have contrasting impacts.

However, ORA is prone to false positives (17). To corroborate these results, replications using ShinyGO and g:Profiler, two commonly used overrepresentation pipelines using different algorithms, were performed next. ShinyGO revealed extensive overlap with WebGestalt for ASO 3+4, but only 5 FDR-significant hits were identified for ASO7+8, although all were also significant by WebGestalt. By contrast, g:Profiler replicated only 2 ASO3+4 FDR-significant hits, both identified by ShinyGO and WebGestalt, but identified extensive overlap with ASO 7+8 WebGestalt (25/26) and ShinyGO (5/5) analyses. Examination of this replicated set is consistent with a role of *ENST8800* exon 1 (ASO3+4) and *ENST1743* exon 3 (ASO7+8) variants in cell cycle regulation and transcriptional regulation, respectively (**Fig. 3B**). Unfortunately, no significant terms were identified for ASO5+6 by any of these approaches.

### Gene set enrichment analysis of *ZFAS1*-suppressed cells reveals a shared group of ontologies linked to cell cycle and translation regulation

By design, ORA uses arbitrary fold-change and significance thresholds to identify patterns that match the most impacted transcripts, thus disregarding most of the dataset. By comparison, Gene Set Enrichment Analysis (GSEA) leverages the entire dataset, i.e. without threshold, and uses a cumulative and ranking approach to identify corresponding functional footprints (18). We reasoned that GSEA’s cumulative approach may recognize transcriptome-wide patterns missed by ORA. Moreover, GSEA provides directional information that is often lacking in ORA, and it has been shown to be less prone to noise than ORA (17). GSEA identified FDR-significant Reactome categories in all 3 treatment groups, with ASO7+8 yielding the most hits (330) (**Supplementary Table S3**). By comparison, ASO3+4 and ASO5+6 affected 132 and 50 pathways, respectively. A shared set of 24 pathways, featuring prominently cell cycle and translation regulation, was identified (**Fig. 4**). Examination of this group revealed, unexpectedly, that ASO3+4 and ASO5+6, both targeting *ENST8800*, had systematically antagonistic effects: whereas ASO3+4 was associated with increased translation-related terms but reduced mitosis terms, the ASO5+6 treatment had opposite effects. By comparison, ASO7+8 suppression was predicted to reduce the activation of these pathways, thus exhibiting a pattern similar to ASO3+4 with regard to the cell cycle but opposite *vis-à-vis* translation-related terms. Moreover, 2 respiration-related terms were reduced in ASO5 +6 and ASO7+8 treated samples but increased in samples treated with ASO3+4.

**Figure 4.**
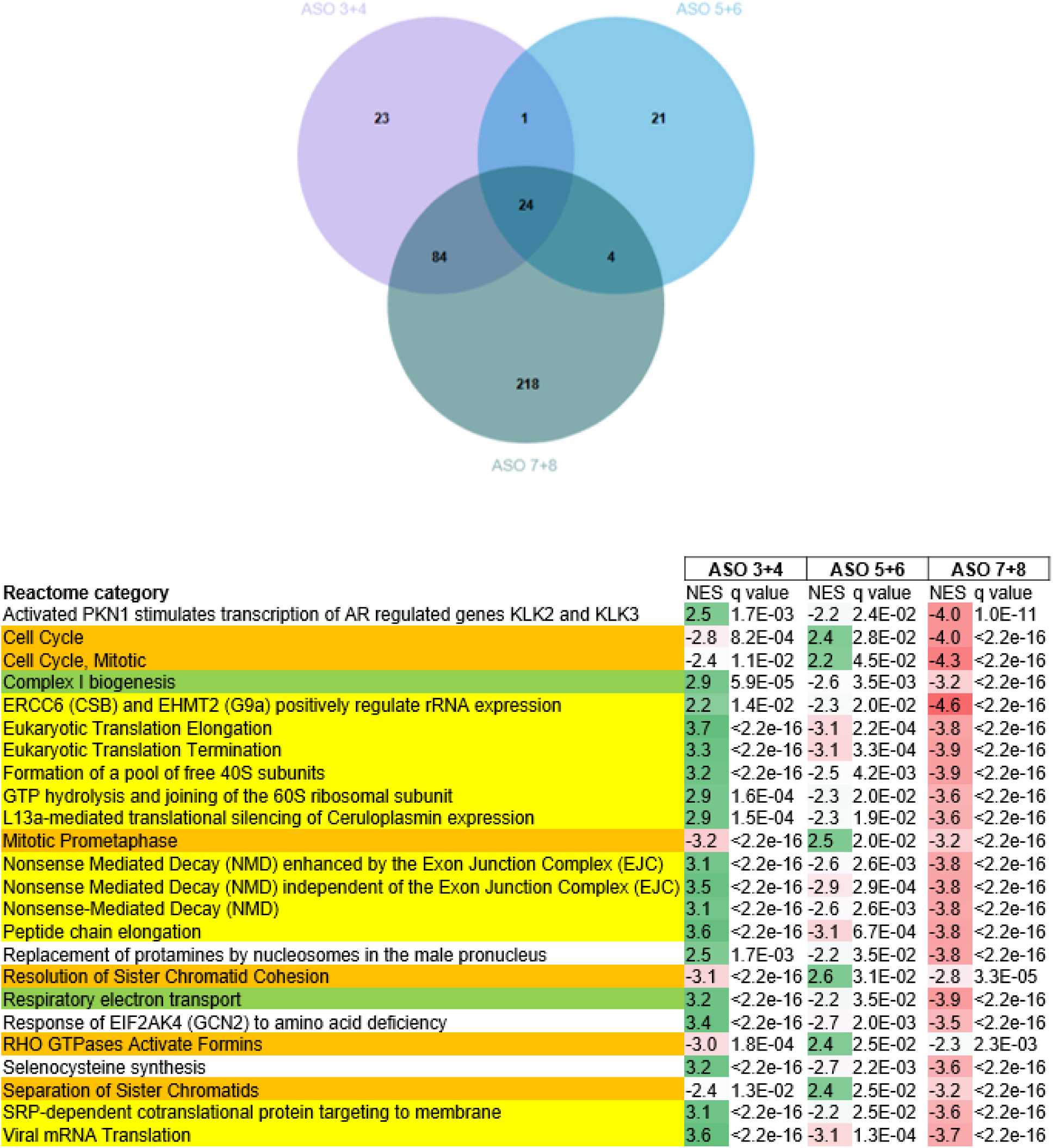
Shared FDR-significant Reactome pathways. Top, Venn diagram of FDR-significant Reactome and Gene Ontology terms identified by GSEA of ASO-treated HepG2 cells. Bottom, shared group of Reactome terms. Terms shared by all three ASO treatments are shown. Terms related to the cell cycle and/or chromosome are highlighted in orange, terms related to translation are highlighted in yellow, and terms related to respiration are highlighted in green. Normalized effect sizes (NES) are color-coded for activation (green) and inhibition (red).

Reactome terms are by definition pathway-centric. To obtain complementary biological information, transcripts were then mapped by GSEA to Biological Process Gene Ontologies (GO), revealing broad sets of enriched ontologies (**Supplementary Table S4**). Similarity clustering of GO categories with REVIGO identified networks of ontologies in each targeted population (**Fig. 5A**). ASO3+4 treatment resulted in a network of increased metabolic capacity, but impaired RNA handling and DNA replication. Similarly, ASO7+8 was predicted to suppress RNA and DNA-related ontologies. However, unlike ASO3+4, ASO7+8 inhibited metabolic terms. By contrast, a single network consisting of increased stress-related terms was identified following ASO5+6 treatment. Comparison of ontologies revealed less inter-ASO overlap than with Reactome terms. Shared terms were, however, concordant with the corresponding Reactome terms. For instance, although only 2 terms were shared by all 3 treatments (chromosome segregation and NADH dehydrogenase complex assembly), chromosome segregation was reduced with ASO3+4/7+8 but increased with ASO5+6, in line with chromatid-related Reactome pathways (**Fig. 5B)**. In summary, although GO analysis identified less shared FDR-significant terms, results were consistent with Reactome enrichment.

**Figure 5.**
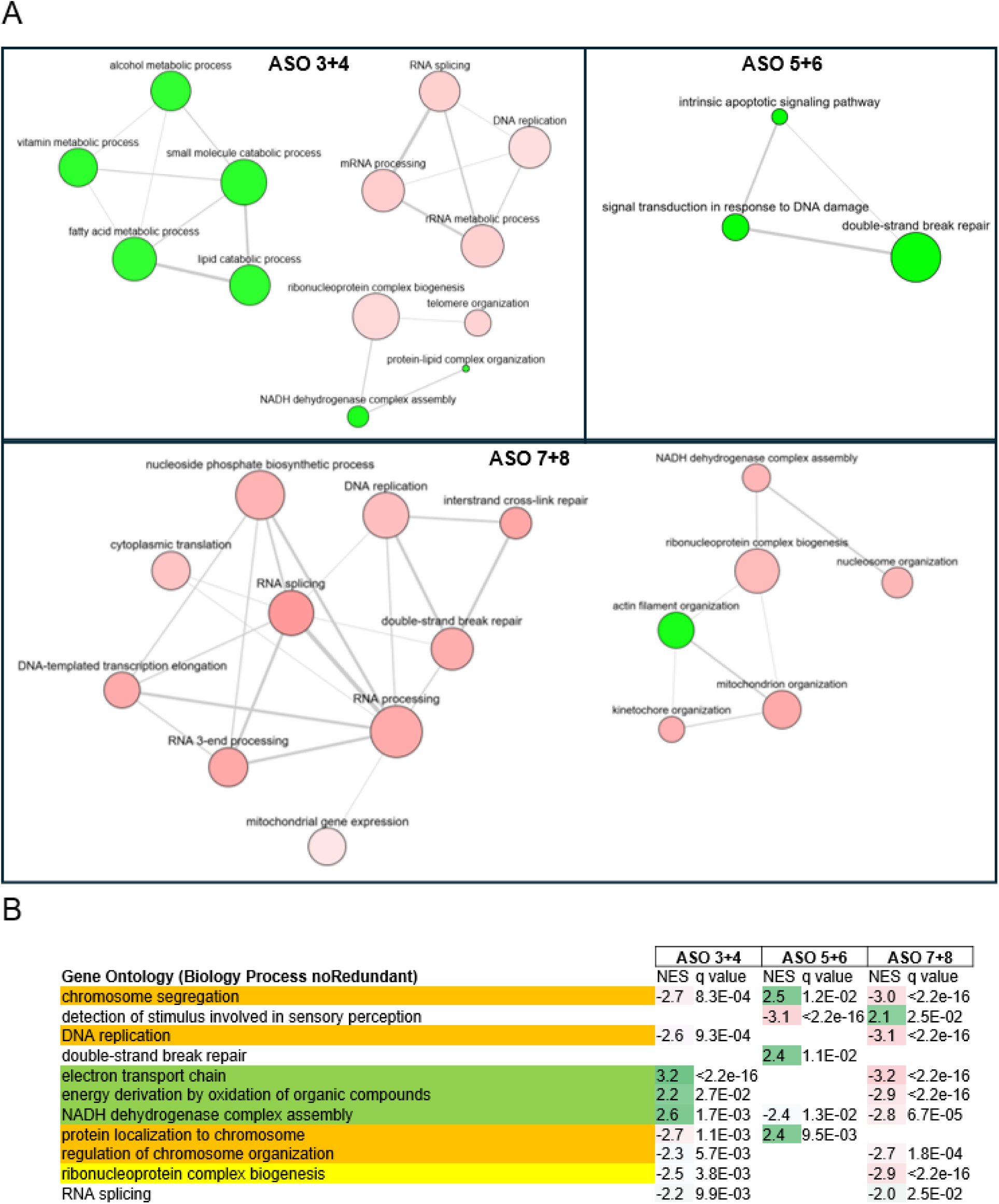
GSEA analysis mapping to GO terms. A, REVIGO clustering of GO terms. Line thickness indicates similarity. Color reflects effect size (negative, red; positive, green). Node size is proportional to the ontology membership. Only groups of >3 nodes are shown. B, Shared Gene Ontology terms. Ontologies shared by two or more ASOs are shown. Terms related to the cell cycle and/or chromosome are highlighted in orange, terms related to translation are highlighted in yellow, and terms related to respiration are highlighted in green. Normalized effect sizes (NES) are color-coded for activation (green) and inhibition (red).

## Discussion

Determining the role of lncRNAs is challenging, primarily due to the absence of a readily accessible code (such as mRNA codons) and poor sequence conservation. Previous studies on *ZFAS1* have focused on variants harboring exons proximal to *ZNFX1*, its protein-coding gene neighbor. This is typically achieved by targeting a proximal *ZFAS1* exon with siRNA or antisense approaches, and/or overexpressing *ZFAS1*, for which relevant isoform or exon information is not systematically provided (e.g., (5,7,19,20)) These studies, for simplicity, ignore putative ZFAS1 “distal” exon-containing isoforms, which may lead to incorrect or incomplete conclusions. For instance, although higher *ZFAS1* expression has been reported in various cancers and is associated with a poor prognosis, the splice variants implicated are poorly defined.

Targeting the abundant *ENST1743* exon 3 (ASO7+8) had a globally suppressive effect, suggesting that the more abundant *ZFAS1* form has a role in promoting proliferation in HepG2 cells, in line with our previous observations (10). However, we provide evidence that other *ZFAS1* splice variants may have distinct, sometimes antagonistic roles, depending on their exon arrangement. For instance, targeting *ENST8800* exon 1 (ASO3+4) and *ENST1743* exon 3 (ASO7+8) had contrasting effects on energy/mitochondria and translation-related terms, whereas both interventions were predicted to reduce cell cycle-related terms by GSEA. These roles would be in line with our previous observation that ZFAS1 variants were predominantly cytosolic (10). This finding supports previous work suggesting that *ZFAS1* facilitates ribosomal maturation in cancer cells, but it also suggests that this effect may be splice variant-dependent (6). Indeed, targeting *ENST8800* exon 1 and *ENST8800* exon 2 had systematically opposite effects on enriched categories, although the impact of *ENST8800* exon 2 targeting was weaker (i.e., lower |NES|). This suggests that both exons may be part of distinct transcripts, exerting contrasting influences. This model is further supported by the suppression of *ENST1743* exon 3, which increased the signal of *ENST8800* exon 2 but not that of *ENST8800* exon 1, as well as the upregulation of *ENST8800* exon 2, which increased *ENST8800* exon 1 abundance. Yet low abundance products spanning *ENST1743* exon 3 and *ENST8800* exon 2, as well as ENST8800 exons 1 and 2, could be amplified by qRT-PCR, indicating that these exons can co-occur in the same transcript, at least occasionally (**Fig. 2E** and (10)). Thus, *ZFAS1* joins a growing list of lncRNAs characterized by multiple splice variants exerting contrasting contributions (21).

Interestingly, although the impact of ASO7+8 was systematically larger on cell cycle terms, these terms were not enriched in the corresponding ORA. However, the transcripts driving most of the enriched ASO7+8 terms were strongly enriched in histones, which have roles above and beyond the identified pathways. For instance, two enriched terms, “Transcriptional regulation of granulopoiesis” and “Defective pyroptosis” (**Fig. 3B**), although seemingly dissimilar, are both driven by the same 11 (out of 16 and 14 genes, respectively) histone transcripts, which were also suppressed with ASO3+4 (although only one reached the expression threshold criteria). Intriguingly, these histone transcripts belong to histone clusters 1 and 2, a group of histones that depend on DNA replication for their expression (22). Thus, ORA may have underestimated the cell cycle impacts of ASO 7+8 suppression.

An intriguing finding relates to the lack of measurable reduction upon targeting *ENST8800* with cognate ASO. Although this may reflect ineffective suppression, overall consistency in the dataset (directionally concordant sets) and reciprocal impacts on enriched categories argue otherwise. High noise, possibly due to the very low expression of *ENST8800* forms, might mask genuine suppression. Alternatively, an inactive *ENST8800*-ASO duplex may form but elude degradation by RNase H. How *ENST8800*, but not *ENST1743*, may escape degradation is unclear, given that the ASOs were designed similarly, incorporating flanking 2′-O-methoxyethyl groups and a phosphorothioate backbone to allow cleavage of the ASO-RNA duplex by RNAse H1 (23). The answer may reside in the insufficient abundance of *ENST8800* to enable the detection and degradation of ASO-RNA heteroduplex by RNase H1. A low-abundance RNA, present at a few copies per cell, would be expected to reach the pM range (assuming a 2000 μm^3^ cell volume), far less than the predicted Kd of RNAse H1 (∼ 1 μM) (24). This finding is reminiscent of our previous observation that the low abundance lncRNA *TRIBAL* was functionally impaired upon cognate ASO treatment, despite inconsistent suppression (25). Thus, we speculate that low-abundance RNA-gapmer complexes may evade degradation and persist as dysfunctional complexes.

We acknowledge limitations of this study. First, the dataset is entirely transcriptomic. As such, the enriched pathways and ontologies were not directly examined. Second, this work examined a single early time point (48 h) to identify proximal regulatory roles. As longer suppression time points were not evaluated, putative late *ZFAS1* contributions might not have been captured. Third, because we have employed an exon-centric approach, the contributions of specific splice variants could not be resolved and will require comprehensive characterization of the *ZFAS1* transcript repertoire. Fourth, as suppression was performed in a transformed human hepatoblastoma cell line, generalizability of these findings to other cell types or species remains to be tested. Lastly, while some off-target effects are expected with antisense interventions, the consistent directionality of the shared differentially affected transcripts and the coherence of enriched categories identified by GSEA support specify, offering new functional insight into *ZFAS1*.

## Supporting information

Supplementary sequence information

## Declarations

### Availability of data and materials

The expression dataset analyzed was deposited at the gene expression omnibus (GEO) and can be obtained from https://www.ncbi.nlm.nih.gov/geo/query/acc.cgi?acc=GSE308839.

## Authors’ contributions

SS designed, performed, analyzed experiments, and wrote the manuscript. PL provided technical assistance, and RM acquired funding and edited the manuscript.

## Acknowledgements

None

## Additional information

### Supplementary Information

**Supplementary Tables.xlsx**. Supplementary Tables S1-4.

**Supplementary Sequence Information.docx**. Oligonucleotide sequences.

