## Supplementary sequence information for "Exon-Resolved Dissection of Shared, Unique, and Antagonistic Functions of ZFAS1 Isoforms"

### Oligonucleotide sequences

#### qPCR primer pairs (forward, reverse)

PPIA
ACCGTGTTCTTCGACATTGC
TTCTGTGAAAGCAGGAACCC

ENST8800ex1
GGTTTTCACAGATCGCTCTGGT
CTCAATTGATCGTAGGGAGCAAAGG

ENST8800ex2
GGAAGTGACAATGTCAGTAGCCC
AGAAGTGCTGGCTGCAGTGT

ENST1743ex3
GTTTATTGTTGGCCAAAACCAGG
TATTCTACTTCCAACACCCGC

ENST8800ex1-ex2
TTGAGATGCAACCAGAGCGAT
CAAGATGAAGGCCTCTTGCC

ENST1743ex3-ex4
TTAGAGCAGCCAGCGGGTA
CTGGTTCAATCAAAGCCTGGTT

ENST1743ex3-ENST8800ex2
TTAGAGCAGCCAGCGGGTA
CAAGATGAAGGCCTCTTGCC

#### ASO gapmers (top, as ordered from IDT; bottom, corresponding nucleotide sequence)

ASOs are 2'-O-(2-methoxy) ethyl-modified oligonucleotides (10 nucleotides; 5 on either side) with a phosphorothioate backbone. Obtained from Integrated DNA Technologies (IDT).

CTLASO

/52MOErC/*/i2MOErC/*/i2MOErT/*/i2MOErT/*/i2MOErC/*C*C*T*G*A*A*G*G*T*T*/i2MOErC/*/i2MOErC/*/i2MOErT/*/i2MOErC/*/32MOErC/
CCTTCCCTGAAGGTTCCTCC

ASO7

/52MOErG/*/i2MOErG/*/i2MOErT/*/i2MOErT/*/i2MOErC/*A*A*T*C*A*A*A*G*C*C*/i2MOErT/*/i2MOErG/*/i2MOErG/*/i2MOErT/*/32MOErT/
GGTTCAATCAAAGCCTGGTT

ASO8

/52MOErC/*/i2MOErT/*/i2MOErT/*/i2MOErC/*/i2MOErC/*A*A*C*A*C*C*C*G*C*A*/i2MOErT/*/i2MOErT/*/i2MOErC/*/i2MOErA/*/32MOErT/
CTTCCAACACCCGCATTCAT

ASO3

/52MOErG/*/i2MOErT/*/i2MOErT/*/i2MOErG/*/i2MOErC/*C*A*A*T*A*C*C*T*G*G*/i2MOErG/*/i2MOErA/*/i2MOErA/*/i2MOErG/*/32MOErG
GTTGCCAATACCTGGGAAGG

ASO4

/52MOErG/*/i2MOErA/*/i2MOErT/*/i2MOErC/*/i2MOErG/*T*A*G*G*G*A*G*C*A*A*/i2MOErA/*/i2MOErG/*/i2MOErG/*/i2MOErC/*/32MOErT
GATCGTAGGGAGCAAAGGCT

ASO5

/52MOErG/*/i2MOErA/*/i2MOErC/*/i2MOErG/*/i2MOErG/*A*C*T*T*G*T*A*C*T*T*/i2MOErC/*/i2MOErC/*/i2MOErC/*/i2MOErT/*/32MOErC
GACGGACTTGTACTTCCCTC

ASO6

/52MOErA/*/i2MOErA/*/i2MOErG/*/i2MOErT/*/i2MOErG/*A*C*A*A*T*G*T*C*A*G*/i2MOErT/*/i2MOErA/*/i2MOErG/*/i2MOErC/*/32MOErC
AAGTGACAATGTCAGTAGCC
